# A phenotype-structured PDE framework for investigating the role of hypoxic memory on tumor invasion under cyclic hypoxia

**DOI:** 10.1101/2025.10.26.684650

**Authors:** Gopinath Sadhu, Paras Jain, Jason Thomas George, Mohit Kumar Jolly

## Abstract

Tumor growth and angiogenesis drive complex spatiotemporal variation in micro-environmental oxygen levels. Previous experimental studies have observed that cancer cells exposed to chronic hypoxia retained a phenotype characterized by enhanced migration and reduced proliferation, even after being shifted to normoxic conditions, a phenomenon which we refer to as *hypoxic memory*. However, because dynamic hypoxia and related hypoxic memory effects are challenging to measure experimentally, our understanding of their implications in tumor invasion is quite limited. Here, we propose a novel phenotype-structured partial differential equation modeling framework to elucidate the effects of hypoxic memory on tumor invasion along one spatial dimension in a cyclically varying hypoxic environment. We incorporated hypoxic memory by including time-dependent changes in hypoxic-to-normoxic phenotype transition rate upon continued exposure to hypoxic conditions. Our model simulations demonstrate that hypoxic memory significantly enhances tumor invasion without necessarily reducing tumor volume. This enhanced invasion was sensitive to the induction rate of hypoxic memory, but not the dilution rate. Further, shorter periods of cyclic hypoxia contributed to a more heterogeneous profile of hypoxic memory in the population, with the tumor front dominated by hypoxic cells that exhibited stronger memory. Overall, our model highlighted the complex interplay between hypoxic memory and cyclic hypoxia in shaping heterogeneous tumor invasion patterns.

## 1 Introduction

Tumor cells require a constant supply of oxygen and nutrients to maintain their rapid growth [1]. In the case of avascular tumors, nutrients diffuse from neighboring blood vessels, which limits the tumor’s size, and also establishes an oxygen gradient inside the tumor region [2] that establishes oxygen-rich (normoxic) and oxygen-deprived (hypoxic) regions. It is generally observed that cells are less proliferative in hypoxic conditions than normoxic ones [3]. However, hypoxic cells are shown to be highly migratory in nature and significantly contribute to tumor invasion [4]. The dichotomy of proliferation and invasion of cancer cells under normoxic and hypoxic environmental conditions has led to the formulation of the ‘go-or-grow hypothesis’ [5, 6]. Numerous experimental studies have demonstrated that the oxygen levels within tumors fluctuate in a spatiotemporal manner over a wide range of timescales, from hours to days, a phenomenon referred to as *cyclic hypoxia* [7, 8]. Cyclic hypoxia arises primarily from the chaotic and leaky vascular network in tumors, which drives variations in oxygen availability [9]. Empirical work replicating such cyclic hypoxic conditions *in vitro* have shown the resulting phenotypic adaptation of cells between normoxic and hypoxic phenotypic states, which affect population growth and migration [10, 11]. However, how different fluctuation characteristics of environmental oxygen would affect the normoxic-hypoxic phenotypic structure of the population and tumor invasion dynamics is less explored, largely because of the experimental limitations. Although prior analyses have helped elucidate how dynamics changes in vasculature and oxygen levels promote tumor progression in general, a specific study of the impact of cyclic hypoxia on tumor invasion still needs to be done [12–14].

In addition to phenotypic adaptation resulting from a fluctuating oxygen environment, hypoxic cells can continue to show enhanced migration even upon re-oxygenation [15]. Such observations suggest that cellular adaptation proceeds based on a ‘memory’ of prior hypoxia. Hypoxic memory is shown to enhance the metastatic ability of cancer cells, with post-hypoxic cells (cells that have experienced hypoxic environment in the past) migrating close to vasculature (normoxic/oxygen-rich regions), and leaving tumor boundary and invading ECM [15–17]. To capture hypoxic memory and explain the distant migration ability of post-hypoxic cells, Rocha et al. [18] hypothesized a persistence time for which post-hypoxic cells remain migratory after returning in normoxic region [19]. However, the timescales at which such memory accrues under hypoxia and diminishes upon re-oxygenation and their subsequent effects on tumor invasion are yet to be investigated.

In this work, we propose a novel phenotype structured partial differential equation (PS-PDE) model to elucidate the effects of phenotypic adaptation during cyclic hypoxia and the associated dynamics of hypoxic memory on tumor invasion in periodic normoxia-hypoxia environmental shifts. The proposed model consists of a set of differential equations that describes the spatio-temporal dynamics of normoxic and hypoxic cancer cells, ECM density, and oxygen concentration. Phenotypic adaptation between normoxic and hypoxic cell states in our model depends on the threshold oxygen level, above and below which cells switch states. Further, hypoxic memory is accounted for by modeling hypoxic cell density as a function of a phenotypic variable controlling the hypoxic-to-normoxic transition rate, in addition to space and time. In this approach, drift in the phenotypic space during hypoxic and normoxic environmental conditions determines the transition rate out of hypoxia. To solve this model, we employ the method of lines (MOL) framework. Simulations reveal that although hypoxic memory resulted in a reduction of tumor volume with respect to the dynamics obtained for the memoryless case, it significantly increased tumor invasion. However, the above trade-off between tumor volume and invasion becomes less pronounced for reduced duration of hypoxic exposure in a periodic cycle. We further observed that tumor invasion is highly sensitive to the rate of hypoxic memory induction, but not to the memory erasure rate. Lastly, our analysis showed that shorter periods of cyclic hypoxia along with slow memory induction timescales led to a varied hypoxic memory profile in the tumor, with the tumor front being primarily composed of hypoxic cells with an established hypoxic memory.

The remainder of the paper is organized as follows. The mathematical model is derived and cast in dimensionless form in Section 2. Section 3 provides a detailed description of the numerical method used to solve the model. Numerical results are presented in Section 4. The paper concludes in Section 5, where we discuss our findings and identify possible directions for future work.

## 2 Model development

In this section, we present a new model of the tumor invasion process occurring in a cyclic hypoxia environment, which accounts for the hypoxic memory of hypoxic cells. We assume that the tumor population consists of two kind of cells, normoxic and hypoxic cells, and is surrounded by an extracellular matrix (ECM). To develop the model, we assume the following underlying biological assumptions:

- Oxygen-dependent cell proliferation and death.
- Depending on the local oxygen concentration, we have hypoxic and normoxic environments and the cells correspondingly attain hypoxic and normoxic phenotypes.
- Both normoxic and hypoxic cells die when oxygen level falls below a fixed value.
- Go-or-grow hypothesis: only hypoxic cells are migratory, and normoxic cells are proliferative in nature.
- Tumor cells employ many mechanisms to degrade the surrounding ECM. In this work, we assume that ECM is collectively degraded by hypoxic tumor cells.
- Duration of hypoxic environments exposure modulates phenotypic switch from hypoxic to normoxic phenotype. Specifically, we consider that the rate of transition from a hypoxic to normoxic phenotype depends on duration of hypoxic environment exposure.

To account for the above desired biological assumptions, we define densities for normoxic cells *n*(**x**, *t*), hypoxic cells *h*(**x**, *t, μ*_*hn*_), ECM *e*(**x**, *t*) and oxygen *c*(**x**, *t*), with **x** = (*x, y, z*) denoting the spatial coordinate, *t* denoting time and *μ*_*hn*_ representing phenotypic space *P* = [0, *μ*_*hn*0_], where *μ*_*hn*0_ is the maximum transition rate of hypoxic cells to normoxic cells under normoxic conditions. The total density of all hypoxic cells in the physical domain is given by

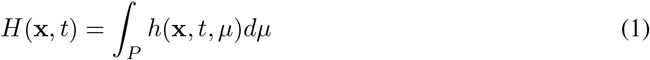

**Normoxic cell dynamics:** The temporal evolution of the normoxic cell density is governed by three primary processes: (i) proliferation, (ii) necrotic death, and (iii) phenotypic switching between normoxic and hypoxic phenotypic states. Proliferation is subject to a volumetric constraint arising from the collective densities of cancer cells and ECM, reflecting spatial limitations within the tumor and its microenvironment. Accordingly, the governing dynamics is expressed as:

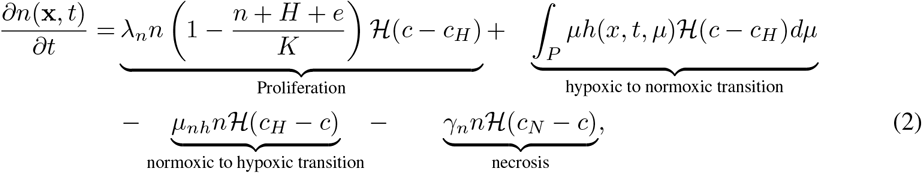

where ℋ is the Heaviside function, defined as

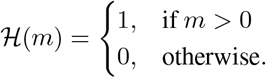

The first term on the right-hand side (RHS) of the Eq. (2) describes normoxic cells that proliferate logistically at a rate of *λ*_*n*_ in a microenvironment with carrying capacity *K* when the oxygen concentration is above the hypoxic threshold value *c*_*H*_. The second term on the RHS indicates that hypoxic cells switch their phenotype to a normoxic one at a rate of *μ*_*hn*_ when the local oxygen concentration exceeds *c*_*H*_. The third term on the RHS defines normoxic cells that adopt a hypoxic phenotype at a rate of *μ*_*nh*_ when the local oxygen concentration drops below *c*_*H*_. Finally, the last term on the RHS represents normoxic cell death at a rate of *γ*_*n*_ when the oxygen concentration falls below the necrotic threshold value *c*_*N*_.

**Hypoxic cell dynamics:** The spatio-temporal dynamics of hypoxic cells is given as

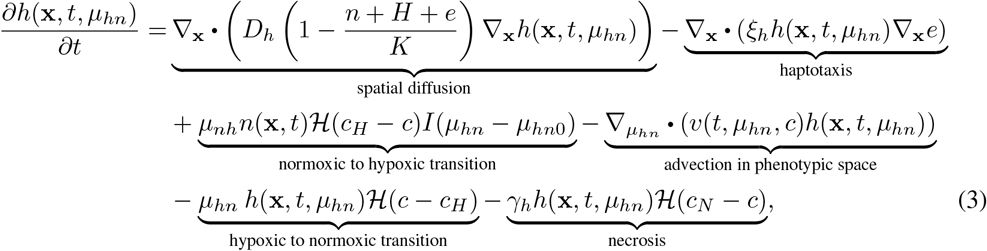

The first term on the RHS of Eq. (3) defines hypoxic cell undirected movement (i.e., diffusion) in the physical space, where this movement is prevented by the presence of hypoxic cells, normoxic cells, and ECM at the current spatial location. The second term on RHS represents hypoxic cells migrating towards high ECM density sites with a rate *ξ*_*h*_, which captures the phenomenon of tumor cells migrating directly towards the region where high adhesive molecules are concentrated (i.e. haptotaxis). The third term on RHS describes normoxic cells adopt hypoxic phenotype with a rate *μ*_*nh*_ when oxygen concentration falls below *c*_*H*_. Here, to account for the build-up of hypoxic memory, we consider that all normoxic cells first switch to the hypoxic state having the highest reverse transition rate *μ*_*hn*_ = *μ*_*hn*0_ upon future exposure to normoxia. This is formulated using the indicator function *I*(*μ*_*hn*_ − *μ*_*hn*0_), defined as follows:

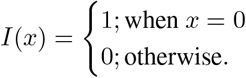

On continued exposure to hypoxia, hypoxic cells density then flows to lower *μ*_*hn*_ phenotypic values, which reduces reverse transitions into normoxic states upon future re-exposure to normoxia. The fourth term highlights this drift of hypoxic cell density in the *μ*_*hn*_ space using the advection term with velocity *v*, defined as follows:

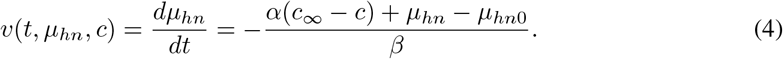

This equation is structured in such a way that the transition rate decreases when the local oxygen concentration in the tumor decreases with respect to its maximum value *c*_∞_. Here, *α* modulates the asymptotic reduction in the transition rate due to hypoxia memory. By setting *α* = 0, we can eliminate the oxygen dependency from *μ*_*hn*_ altogether, resulting in a constant value *μ*_*hn*_(*t*) = *μ*_*hn*0_, which represents the absence of hypoxic memory in our modeling framework. To analyze hypoxic memory cases, we set *α* such that minimal positive *μ*_*hn*_ is asymptotically attained on continued hypoxia exposure. The parameter *β* regulates the rate at which *μ*_*hn*_ changes, and can be set according to the experimental observed timescales of hypoxia memory induction and withdrawal (Appendix 6.3). The fifth term describes hypoxic to normoxic state transition at the rate of *μ*_*hn*_ when oxygen concentration rises above *c*_*H*_. The last term defines hypoxic cell death at a rate *γ*_*h*_ when the local oxygen level falls below *c*_*N*_.

**ECM dynamics:** Here, we have considered that the ECM is collectively degraded by all phenotypic hypoxic cells in the physical space (given by Eq. 1) [20] with a rate *δ*. Hence, the governing equation for ECM is given as

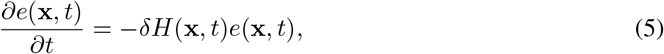

In this work, we consider oxygen field in tumor microenvironment is dynamics with time, which mimics *in vitro* protocol of cyclic hypoxia. In this protocol, oxygen is spatially homogeneous. This cyclic hypoxia is mathematically formulated as

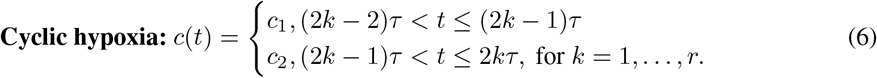

Where 2*rτ* = *T*_*final*_ is total time of interest, 2*τ* is a period of cyclic hypoxia, *c*_1_(*> c*_*H*_) is oxygen-rich environment and *c*_2_(*< c*_*H*_) is hypoxic environment. The typical time frame of cyclic hypoxia is equal to half to five time of cell proliferation time scale i.e., 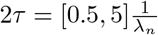.

### 2.1 Nondimensionalization

In order to solve the system numerically, we first of all non-dimensionalise the equations. The variables and parameters in the system equations and their associated boundary conditions are transformed into dimensionless quantities using the following reference variables.

- Reference length scale, *L*, (e.g., the maximum invasion distance of the cancer cells at this early stage of invasion 0.1 − 1 cm),
- reference time unit, 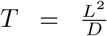, where *D* is a reference chemical diffusion coefficient, e.g., 10^−6^ *cm*^2^*s*^−1^. Therefore, we deduce that *T* varies between 10^4^ − 10^6^ *s*.
- Reference cell density *K*, oxygen concentration *c*_1_

We thus define the non-dimensional variables:

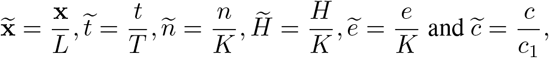

and new parameters via the following scaling: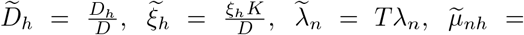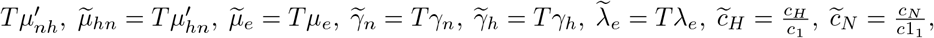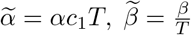

For compactness of notation, we drop the tilde from variables and parameters in the non dimensional model. The dimensionless governing equations can then be written in the following general form:

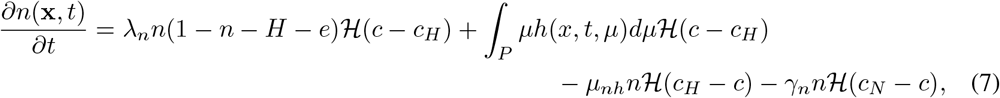

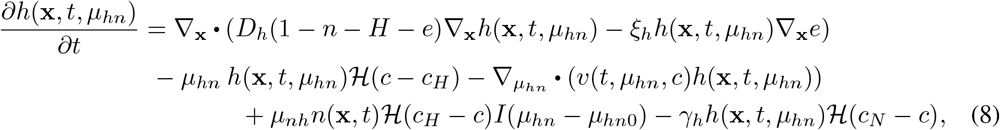

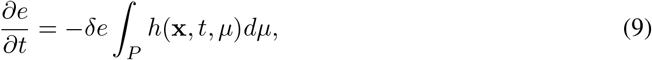

where 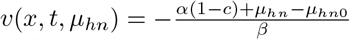.

The non-dimensional form of oxygen field in cyclic hypoxia is given as

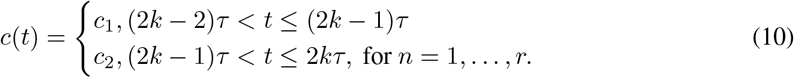

Where 2*rτ* = *T*_*final*_ is total time of interest and 2*τ* is a period of cyclic hypoxia.

### 2.2 Boundary and initial conditions

To close the system, we need to prescribe the appropriate initial and boundary conditions for Eq (7)-(9). In one dimension, the physical domain is [0, *X*] along the *x*-axis. We assume that hypoxic tumor cells do not leave the computational domain [0, *X*] and phenotypic space [0, 1]. The boundary conditions for Eq. (8) are given as

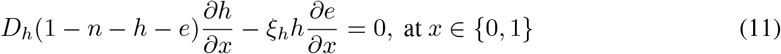

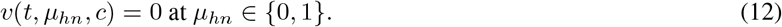

**Initial conditions:** Initially, we consider that tumor consists only normoxic cells and surrounded by the ECM. Hence, the corresponding initial conditions are given in physical domain as

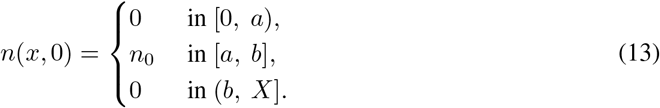

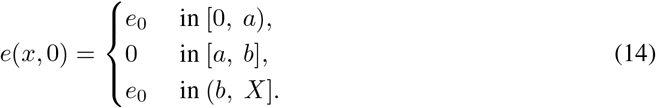

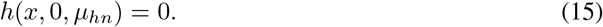

Here, we consider *n*_0_ = 0.5, *e*_0_ = 0.5, *a* = 4, *b* = 6 and *X* = 10.

### 2.3 Model parameter values

The most of parameter values in the model are collected from the existing literature. The remaining parameter values are chosen within suitable regimes such that the results are consistent with the physics of the problem. Table 1 represents the description of physical meaning of parameters and both dimensional and dimensionless values.

**Table 1:** Parameter descriptions and their both dimensional and non-dimensional values.

| Parameter | Description | Dimensional value | Dimensionless value | Source |
| --- | --- | --- | --- | --- |
| $D_h$ | Diffusivity of hypoxic cells | $5 \times 10^{-10} - 2.5 \times 10^{-9} \text{ cm}^2 \text{ s}^{-1}$ | 0.01 | [21] |
| $\lambda_n$ | Proliferation rate of normoxic cells | $0.014 - 0.056 \text{ hr}^{-1}$ | 0.1 | [14] |
| $\mu_{nh}$ | Transition rate from normoxic to hypoxic | $1 \text{ hr}^{-1}$ | 0.5 | [22] |
| $\gamma_n$ | Necrotic death rate of normoxic cells | $\frac{\lambda_n}{10}$ | 0.001 | Assumed |
| $\gamma_h$ | Necrotic death rate of hypoxic cells | $\gamma_n$ | 0.001 | Assumed |
| $\delta$ | ECM degradation rate | — | 15 | [21, 23] |
| $c_H$ | Hypoxic threshold value of oxygen | $5 \text{ mmHg}$ | 0.5 | [14] |
| $c_N$ | Necrotic threshold value of oxygen | $1 \text{ mmHg}$ | 0.1 | [14] |
| $\xi_h$ | Haptotaxis coefficient | — | 0.001 | [21, 23] |
| $\alpha$ | Strength of memory induction | — | 0.8 | Assumed |
| $\beta$ | Timescale of memory induction/erasing | — | 10 | Assumed |
| $\mu_{hno}$ | Basal transition rate of hypoxic to normoxic | $\frac{1}{96} \text{ hr}^{-1}$ | 0.5 | [22] |

## 3 Numerical method

To solve the proposed model, we employed the Method of Lines (MOL) in a finite difference framework. The physical domain [0, *X*] is divided into *N* − 1 intervals of uniform length 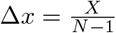, while the phenotypic domain [0, 1] is partitioned into *M* − 1 intervals of uniform length 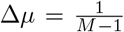. The system of differential equations is discretized explicitly: central difference schemes are applied for spatial derivatives, while forward differences are used for temporal derivatives. The drift (convective) term in Eq. (8) is discretized using a first-order upwind scheme (Eq. (18)).

Let Δ*t* denote the time step, and let computations proceed up to final time *T*_final_, such that *p*Δ*t* = *T*_final_, where *p* is the number of time steps. We define the discrete variables as follows: 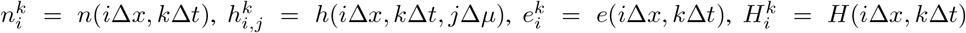, and 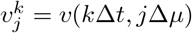 where *i* ∈{0, 1, …, *N* − 1}, *j* ∈ {0, 1, …, *M* − 1}, and *k* ∈ {0, 1, …, *p*}. The difference equation for Eq. (7) is obtained as,

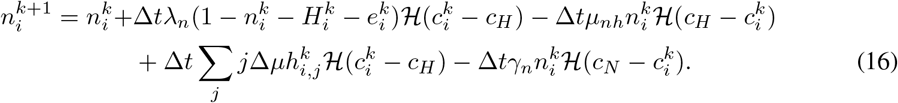

The resulted difference equation for hypoxic cells density, given by Eq. (8), is obtained as,

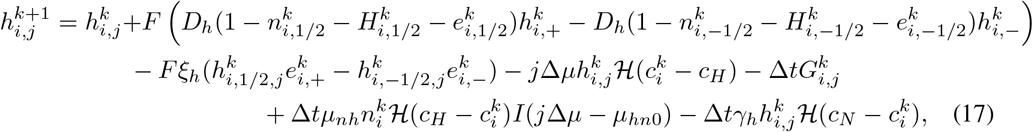

where

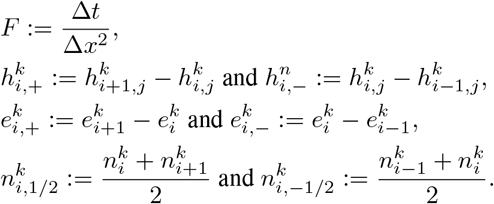

Here, 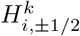 and 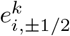 carry similar meaning.

The term 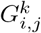 defines convection term at (*i, j*) and it is discretized using first order upwind scheme, given as

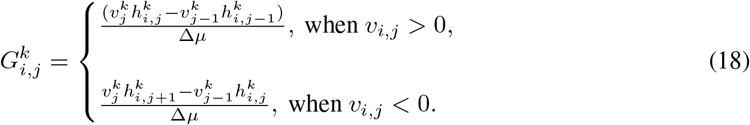

At the boundary nodes, we employ no flux boundary condition at *i* = 0 and *i* = *N* − 1 and zero convective flux at *j* = 0 and *j* = *M* − 1.

The Eq. (9) is discretized as

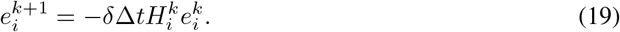

The numerical stability is ensured through the Courant–Friedrichs–Lewy (CFL) condition, which is defined as

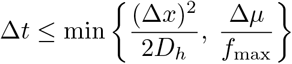

## 4 Results

### 4.1 Spatially homogeneous oxygen field

Here, we have simulated the model Eqs. (7)-(9) with the corresponding boundary and initial conditions, given in Eqs. (11)-(15) in two cases: (i) oxygen-deprivation (*c < c*_*H*_) and (ii) oxygen-rich (*c > c*_*H*_) throughout simulation time.

**Case-I: oxygen-deprivation (*c < c*_*H*_) oxygen field:** We set the oxygen field in the tumor growth medium below the hypoxic threshold, i.e., *c* = 0.4 *< c*_*H*_. Numerical simulations show that normoxic 9 tumor cells gradually convert into hypoxic cells, and the tumor becomes fully hypoxic after *t >* 12 (Figure 1A). Note that these numerical dynamics of total normoxic and hypoxic cell populations strike great agreement with our analytical results in Eq. (20) (the derivations of the analytical expressions are shown in the Appendix 6.1). We also observe that the tumor begins to invade the surrounding ECM as hypoxic cells degrade it (Figure 1B ii). Furthermore, Figure 1C shows that when tumor cells are exposed to a hypoxic environment, normoxic cells adopt a hypoxic phenotype, with their entry starting from *μ*_*hn*_ = 0.5. It is observed that the hypoxic population drifts toward lower values of *μ*_*hn*_ when the tumor is exposed to hypoxia for a prolonged period, indicating the induction of hypoxic memory.

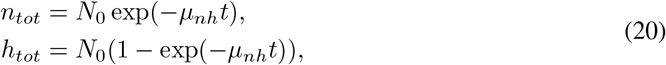

where *n*_*tot*_ and *h*_*tot*_ are total densities of normoxic and hypoxic cells; *N*_0_ is total initial density of normoxic cells.

**Fig. 1:**
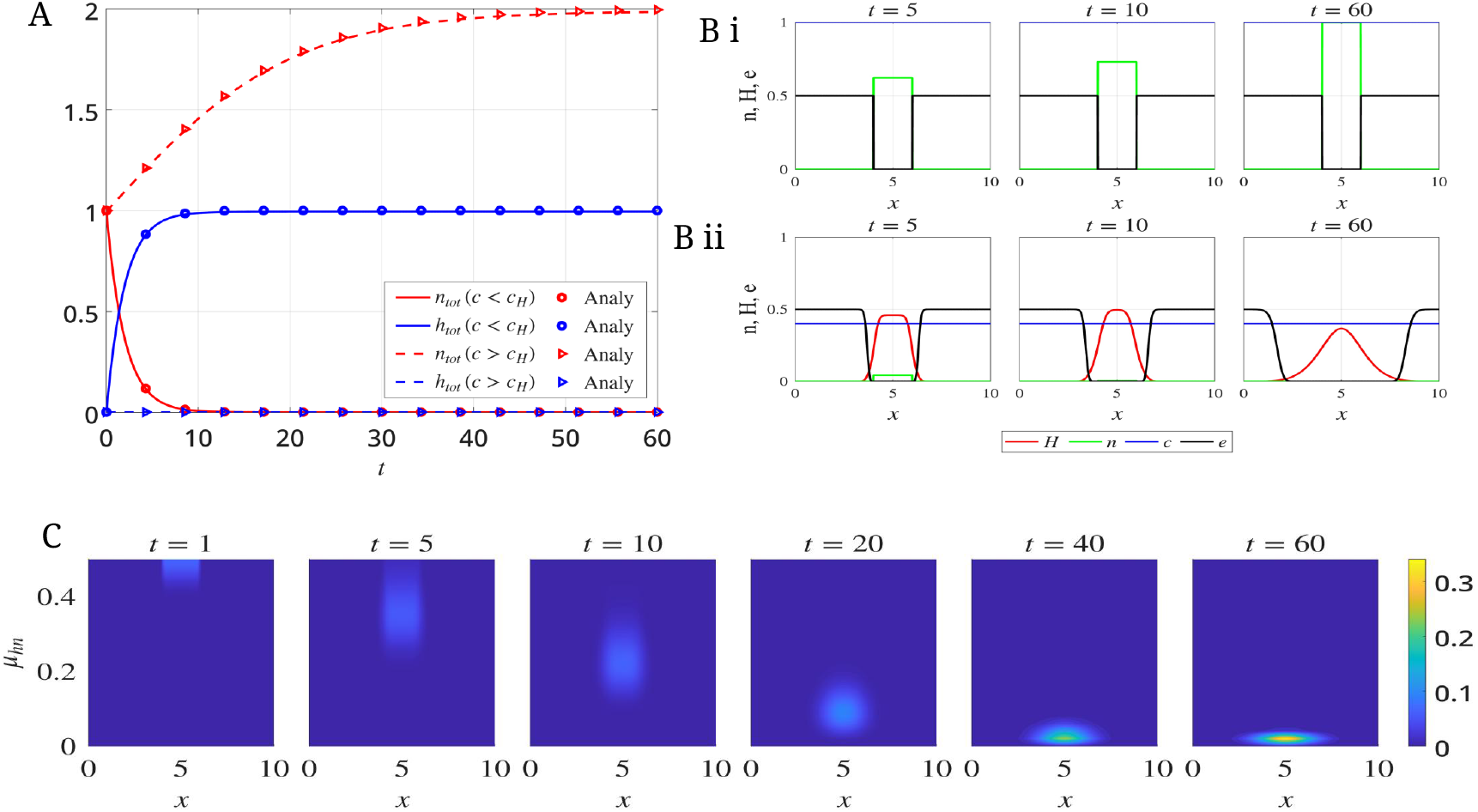
(A) Time evolution of normoxic and hypoxic cell mass in oxygen fields *c* = 0.4 *< c*_*H*_ and *c* = 1 *> c*_*H*_, obtained numerically and analytically. (B) Spatial distributions of hypoxic cells, normoxic cells, ECM density, and oxygen concentration at *t* = 5, 10, and 60 in (i) *c* = 1 *> c*_*H*_ and (ii) *c* = 0.4 *< c*_*H*_. (C) Distribution of hypoxic cell density across the physical and phenotypic domains for *c* = 0.4 *< c*_*H*_.

**Case-II: oxygen-rich (*c > c*_*H*_):** We fix the oxygen concentration in the tumor medium at a level above the hypoxic threshold. Under this condition, the total mass of normoxic cells increases over time (Figure 1A), as normoxic cells proliferate in the oxygen-rich environment. This result agrees closely with the analytical result in Eq. (21), derived in the Appendix 6.2. In addition, we observe that tumor expansion is arrested because normoxic cells do not switch to hypoxic cells, which are responsible for ECM degradation (Figure 1B i). Furthermore, hypoxic memory is not expected to arise, as the cells remain in an oxygen-rich environment.

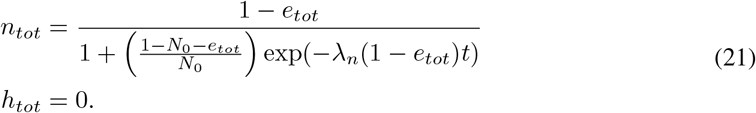

Here, *e*_*tot*_ is the total density of ECM, which is constant in this scenario.

### 4.2 Symmetric duration of normoxic and hypoxic exposure in cyclic hypoxia

In this section, we investigate how periods of cyclic hypoxia influence tumor invasion. The timescale of cyclic hypoxia typically ranges from hours to days [8, 24]. In this study, we select the timescale relative to the proliferation rate, given by 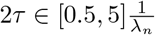. For numerical exploration, we consider 2*τ* = 5, 10, 15, and 20, ensuring that the total durations of exposure to hypoxic and normoxic environments are equal by the end of simulation (t = 60).

The first row of Figure 2A demonstrates that tumors subjected to rapidly cycling hypoxia display highly invasive characteristics. However, such rapid cyclic hypoxia reduces tumor volume (second row of Figure 2A). To understand the above observation, we next examined the dynamics of the hypoxic cell fraction in the population. We found that shorter periods of cyclic hypoxia resulted in higher mean hypoxic cell fractions present in the tumor front relative to longer periods in the simulated time-frame (compare mean hypoxic cell fractions denoted by blue crosses across periods in Figure 2B, Figure S1A). And such increasing share of hypoxic cells leads to higher invasion. Further, since the total duration of normoxic and hypoxic environments in the simulations are same across different time periods, and that the transition rates between normoxic and hypoxic states happen at same rates (*μ*_*nh*_ = *μ*_*hn*0_ = 0.5), we reasoned that the reduced growth of normoxic cells having shorter periods occur as a result of intermittent growth phases and repeated phenotypic adaptation with shorter periods as compared to persistent growth phases with longer periods.

**Fig. 2:**
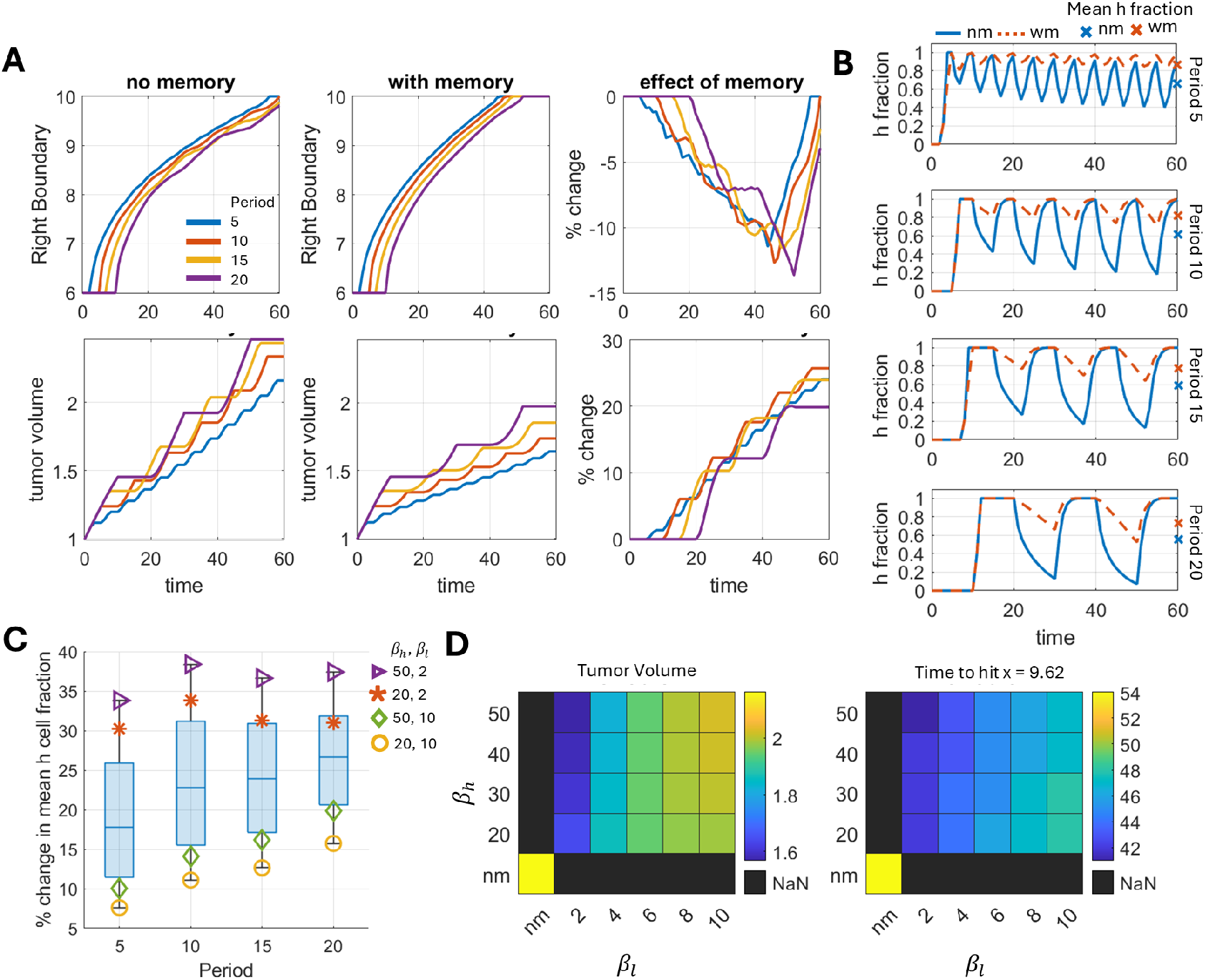
Enhanced tumor invasion in the presence of hypoxic memory. A) The tumor right boundary and tumor volume evolution over time in a cyclic hypoxic environment with variable periods along with cells exhibiting hypoxic memory. B) Time evolution of the hypoxic cell fraction at the tumor front for different value of periods. C) Combined effects of memory erasure (*β*_*h*_) and induction (*β*_*l*_) timescales on the percent change of mean hypoxic cell fraction. D) Effects of *β*_*h*_ and *β*_*l*_ on tumor volume at the end of simulations (*t* = 60), and time to reach (hit) physical space of *x* = 9.62 for period 5. For panels A and B, the time scales of memory induction and memory erasure are *β*_*l*_ = 2, *β* = 20, respectively. In panel B and C, mean hypoxic (h) fractions were quantified by averaging hypoxic cell fractions at the tumor front over the entire simulation duration. Here, the tumor front is defined by spatial region from the tumor right boundary to 0.5 units distance inside, with right boundary being the spatial position *x′* where *n*(*x′, t*) + *H*(*x′, t*) = 10^−6^ for a given *t*.

The above observations on tumor volume and invasion in cyclic hypoxia with no hypoxic memory raises the intriguing question of how hypoxic memory would effect such behaviors. Experimental reports on phenotypic memory of different cellular programs suggest that memory induction occurs on the timescale of cellular division, while memory erasure occurs on the timescale of multiple cell divisions [25–30]. To align with these estimates, we sampled memory erasure timescales from four to ten cell divisions, and memory induction timescales from half to two cell divisions for our analysis by varying *β*_*h*_ ∈ [20, 50] and *β*_*l*_ ∈ [2, 10] (Appendix 6.3). A higher value of *β*_*h*_ indicates a slower rate of decay of hypoxic memory when hypoxic cells are placed in an oxygen – rich environment. Conversely, a lower value of *β*_*l*_ suggests a rapid induction of hypoxic memory when tumor cells are exposed to a hypoxic environment. Simulation results predict that hypoxic memory reduces tumor volume and enhanced migration across all periods of cyclic hypoxia, compared to cases without memory (Figure 2A). It happens because hypoxic memory increases the fraction of hypoxic cells independent of the hypoxia cycling period (Figure 2B). Additionally, we have measured the relative change in tumor volumes and invasion between the memoryless and hypoxic memory cases across simulation time (Figure 2A effect of memory). Our results suggest that hypoxic memory exerts a maximal effect on tumor volume and invasion (right boundary hitting time) for a period of 10 oscillations, which also coincides with the same period that generates maximal differences in mean hypoxic cell fractions between memory and memoryless cases for the given choice of memory induction and dilution timescales - *β*_*l*_ = 2 and *β*_*h*_ = 20 (Figure 2C).

We next performed a series of simulations to investigate the effects of memory erasure and induction timescales on tumor volume and invasion dynamics. We find that while changes in *β*_*h*_ significantly impact tumor volume and invasion speed (measured as hitting times to the right boundary), changes in *β*_*h*_ have minimal impact (Figure 2C). Further, coupling rapid memory induction (*β*_*l*_ = 2) with slow erasure (*β*_*h*_ = 50) slows growth but promotes invasion. In contrast, coupling slow induction (*β*_*l*_ = 10) with fast erasure (*β*_*h*_ = 20) promotes growth but slows invasion (Figure 2D, Figure S1B).

### 4.3 Influences of hypoxic bias on tumor invasion and growth dynamics

Empirical studies have suggested that oxygen oscillations between normoxic and hypoxic levels at the tumor site may not be equal in duration [24, 31]. Therefore, we next examined how hypoxic bias affects tumor growth and invasion dynamics, where hypoxic bias is defined as the fraction of hypoxic environment duration in a periodic cycle. For example, a hypoxic bias of 0.25 represents hypoxia environment to be present for a quarter of the environment period. We performed a series of simulations with bias values of 0.25, 0.4, 0.5, 0.6, and 0.75.

Simulated results with or without hypoxic memory indicated that increasing hypoxic bias reduced tumor volume and enhanced tumor invasion (Figure 3A, B). This trend is because, with increasing hypoxic bias, the tumor experiences shorter exposure to oxygen-rich conditions, which limit its growth but promotes invasion by retaining cells in the hypoxic state Further, in a weak hypoxia bias environmental cycle (Hypoxia bias 0.25), the presence of hypoxia memory significantly elevates tumor invasion with minimal reduction in the tumor volume with respect to memoryless case. Thus, the trade-off between tumor volume and invasion resulting from hypoxic memory can be tuned based on the composition and period of the fluctuating environment (Figure 3B, C).

**Fig. 3:**
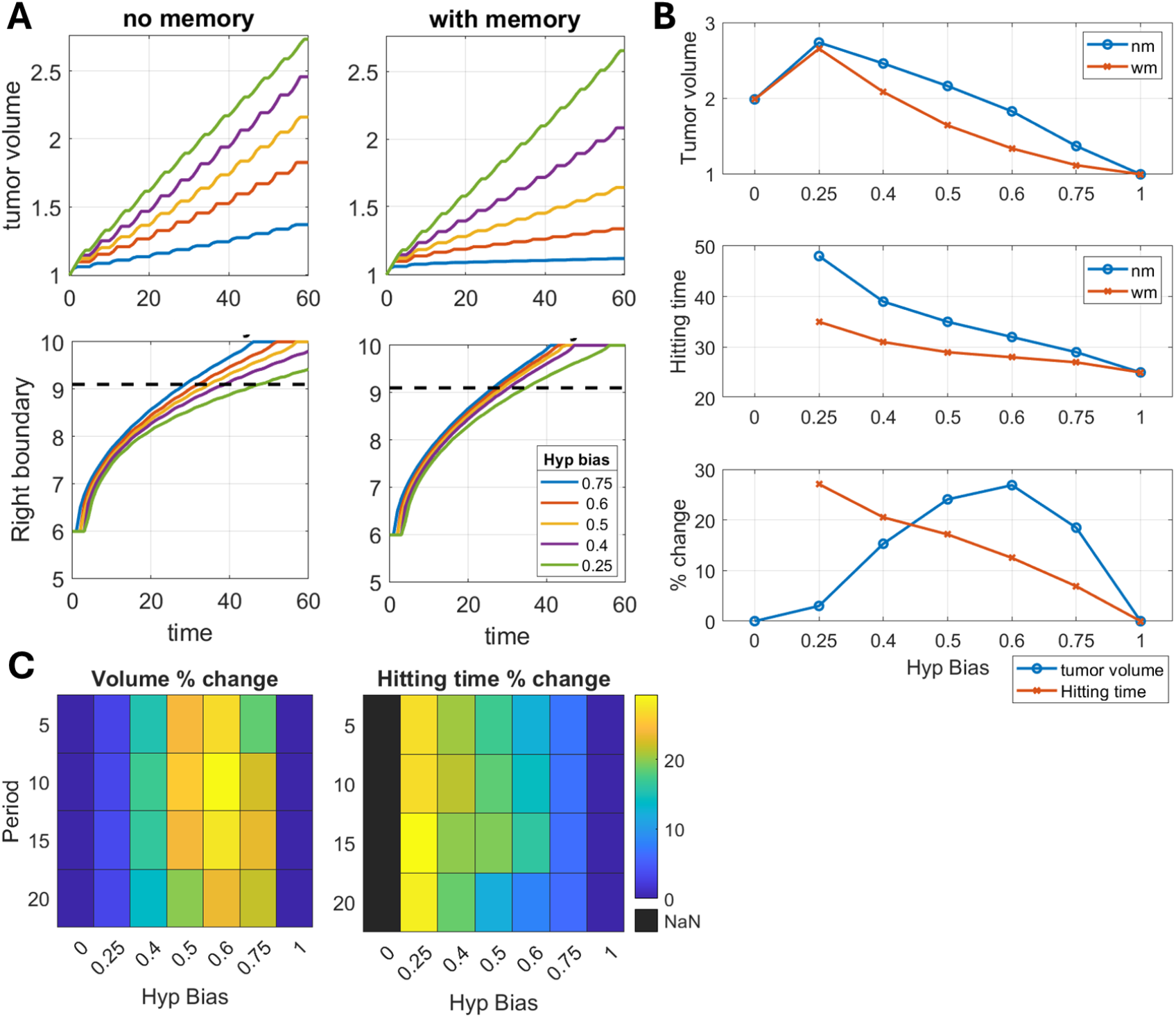
Trade-off between invasion and tumor volume due to hypoxic memory diminishes for shorter duration of hypoxic environment in the periodic cycle. A) Time dynamics of tumor volume and tumor front for different fractions of hypoxia duration (termed as hypoxic bias) in an environmental cycle of period 5. The dynamics is analyzed for the memory and memoryless cases. B) Effects of hypoxic bias of the environmental cycle on tumor volume at the end of simulations (t = 60), and the *x* = 9.1 hitting time for the memory and memoryless cases. Percent change represents change in the tumor volume and hitting times for hypoxic memory cases with respect to corresponding no memory cases. C) Combined effects of hypoxia bias and period of the environmental cycle on the percent change of tumor volume and hitting time due to hypoxic memory with respect to corresponding memoryless cases. For panel A and B environment period is 5. In all panels memory induction rate (*β*_*l*_) is 2 and withdrawal (*β*_*h*_) is 20.

### 4.4 Spatiotemporal changes in memory structure under cyclic hypoxia

So far we have observed the effect of hypoxic memory on tumor volume and invasion. However, the observed collective effect results from dynamics changes in hypoxic memory distribution within hypoxic cells. These changes can be quantified by measuring the distribution of hypoxic cell density in the *μ*_*hn*_ space at two distinct time points – 1) end of a normoxic cycle and 2) end of a hypoxic cycle, and at two distinct spatial locations – 1) bulk and 2) tumor front (spatial region comprised of the tumor right boundary to 0.5 units distance inside). For shorter time period with symmetric hypoxia environmental cycle (hypoxic bias 0.5), we found only minor differences in the memory structure resulting after hypoxia and normoxia environmental cycles, both in the bulk and front spatial regions (Figure 4A - period 5, Figure S2). However, we notice that the front region is dominated by stronger memory cells than the bulk. The differences in the memory structure of bulk and front cells are retained with the variation of the memory induction and withdrawal rates as quantified using Jensen-Shannon Divergence (JSD) between the two phenotypic distributions (Figure S3). The differences in memory structure between bulk and front regions reduce for increasing period of cyclic hypoxia, although the front is still dominated with strong memory cells as compared to the bulk (Figure 4A - period 20, Figure 4B, Figure S4-7). Further, increasing cyclic hypoxia period also lead to differences in the memory structure resulting at the end of hypoxic and normoxic environmental cycles.

**Fig. 4:**
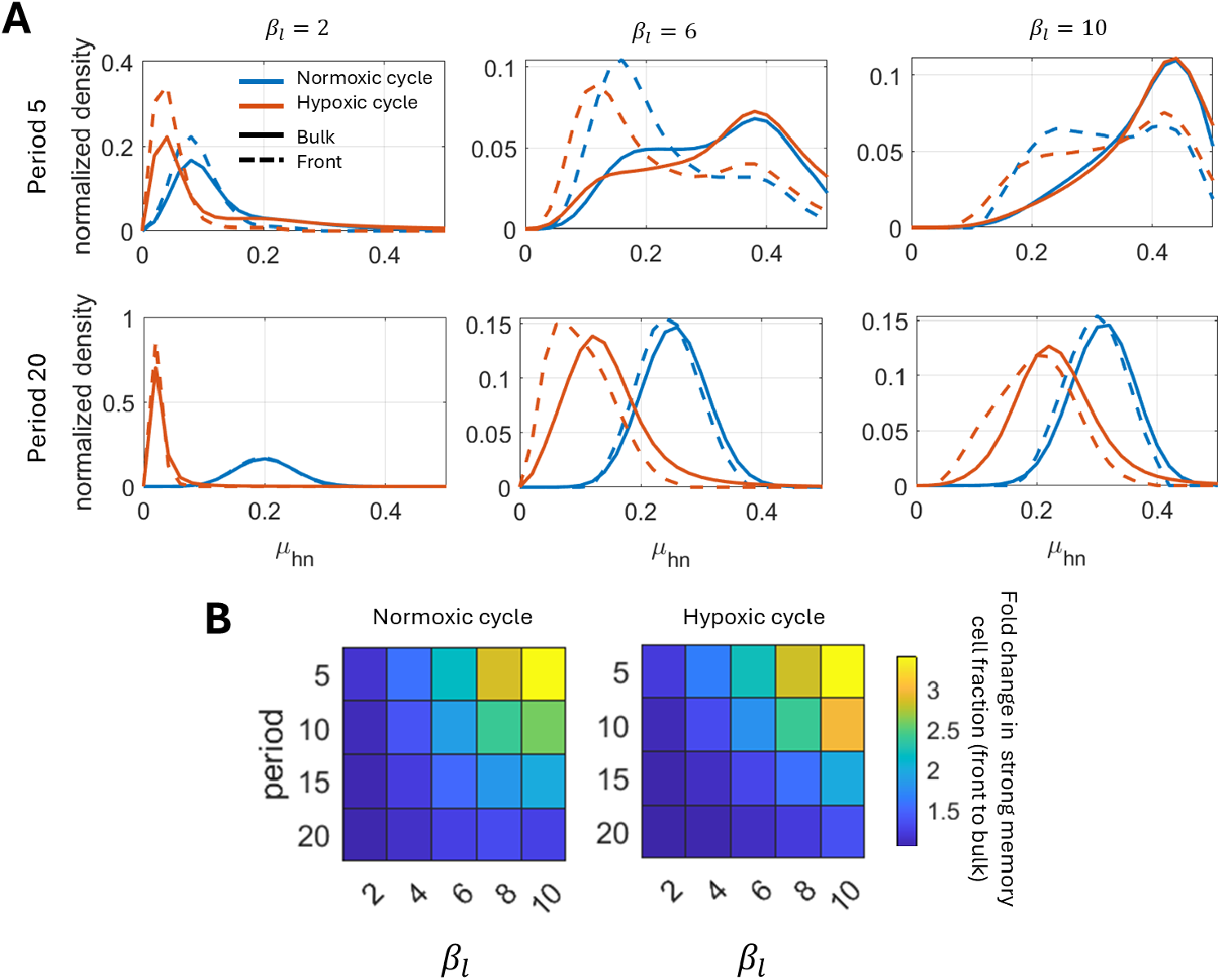
Cells with strong hypoxic memory dominate the invasive front. A) Normalized phenotypic (*μ*_*hn*_) distribution of cells in the bulk (total population) and front (defined as the region within 0.5 units of distance from the right boundary) analyzed at the end of the normoxic and hypoxic cycles of the last period of the simulation. The hypoxic memory induction timescale is varied while memory withdrawal timescales are fixed (see titles of the first row plots). B) Quantification of ratio of strong memory cell fraction in the front to bulk regions at the end of normoxic and hypoxic cycles. Here, the strong memory fraction is calculated by summing normalized cell density with *μ*_*hn*_ *<* 0.25. Results presented in both panel A and B are with *β*_*h*_ = 20 and hypoxia environment bias 0.5. The tumor right boundary is the spatial position *x′* where *n*(*x′, t*) + *H*(*x′, t*) = 10^−6^ for a given *t*.

## 5 Discussion

Intratumoral oxygen concentrations fluctuate spatio-temporally, giving rise to cyclic hypoxia. And, such dynamic environmental changes lead to normoxic-hypoxic phenotypic switching among the cancer cells, contributing to intratumoral heterogeneity [14, 24, 31]. Further, post-hypoxic cells are shown to sustain their phenotype even under normoxic environments — a phenomenon termed as hypoxic memory. Since hypoxic environment can vary over different timescales, hypoxic memory can play a critical role in the invasion process [32, 33], however, this largely remains unexplored. In this study, we propose a novel PS-PDE model to investigate the changes in invasion dynamics under cyclic hypoxia with and without hypoxic memory. Here, the cyclic hypoxia is mimicked by tightly regulating oxygen concentrations between two levels as is typically done in an *in-vitro* setup; and the hypoxic memory is modeled through time-dependent reduction in the hypoxic-to-normoxic transition rates based on continued hypoxia exposure.

Phenotype-Structured Partial Differential Equation (PS-PDE) models have been extensively used to study spatial and non-spatial tumor dynamics over the past decade. This general class of models help to associate population-level behavior with cell state dynamics, capturing time-variant properties of cellular events [34–38]. Ardaševa et al. [39] proposed a non-spatial phenotype-structured model to study the evolutionary cross-talk between population growth and metabolic heterogeneity (spectrum of oxidative phosphorylation - glycolysis states) in fluctuating oxygen environments. Recently, a gene regulatory network was used to inform growth and phenotypic transitions to study the dynamics of epithelial-mesenchymal heterogeneity in a population of cancer cells [40]. To capture the dynamics of tumor progression in a spatial context, Lorenzi and Painter [41] studied the influence of local chemo-attractant densities on the transitions between proliferative and chemotactic phenotypes, which collectively determine the phenotypic structure of the growing population. Similarly, the influence of local cell and ECM densities on transitions between proliferative and invasive phenotypes has also been studied using a discrete-phenotype PDE model [42]. Thus, PS-PDE models are an appropriate framework to couple cellular changes with spatiotemporal dynamics during tumor progression.

As we currently lack detailed molecular-level insights on hypoxic memory, the timescales of hypoxic memory induction and erasure timescales are not yet established. Therefore, for our analysis, we selected ranges of hypoxic memory timescales based on existing knowledge of memory timescales for other well-characterized cellular processes [25–29]. Thus, we considered the timescales of memory dilution to range from four to ten cell divisions (Appendix 6.3), a timescale typical of protein turnover and chromatin-based epigenetic remodeling. Further, since existing experimental reports captured hypoxia memory effects after short-term hypoxia exposure (∼1 generation, [32]), we assumed hypoxic memory induction to happen on much faster timescales, from one-half to two times the duration of cell divisions (Appendix 6.3).

Our simulation results highlighted that the hypoxic cells may not necessarily reduce tumor volume; however, they do affect local invasion significantly. Our findings suggest that a shorter period of cyclic hypoxia enhances the hypoxic cell fraction in the tumor, leading to higher invasion. And, the presence of hypoxia memory further adds to the hypoxic fraction, which significantly elevates invasion but reduces tumor volume, providing reminiscence of observations reported in [15, 43]. On increasing the hypoxic bias of the cyclic environment, we found gradual decrease in tumor volume with concomitant increase in invasion speed. These observations highlight how the balance between hypoxic and normoxic conditions, together with the timescales of hypoxic memory induction, differentially regulates tumor expansion and invasive behavior.

While analyzing the dynamics of hypoxic memory structure, we observed significantly different memory structure at the tumor bulk and front regions for −1) short-vs-long periodic fluctuations, and 2) slow-vs-fast memory induction. And, because of the coupled nature of various cellular processes, the cellular heterogeneity in hypoxia memory can orchestrate as heterogeneity in IFN-signaling, antigen presentation [15], epithelial-mesenchymal heterogeneity [44], and drug resistance [17]. Thus, long-term hypoxia exposure and associated hypoxic memory may drive tumor progression through multiple manifestations than just invasion. As hypoxia can be a major barrier to the clinical success of radiotherapy and chemotherapy [45, 46], it would also be valuable to explore how hypoxic memory influences therapeutic efficacy.

Our work suffers from various limitations. First, it only explains the effects of hypoxic memory on tumor invasion where oxygen levels are spatially homogeneous and vary only with time. However, in spheroid tumors or *in vivo* tumors, oxygen concentrations are not spatially homogeneous [47, 48]. The current study could be extended to investigate the role of hypoxic memory in tumor invasion dynamics under spatiotemporally heterogeneous oxygen fields. Second, it adopts a phenomenological approach without getting into mechanistic details of cellular responses to cyclic vs. chronic hypoxia, which can sometimes be opposite to one another as well [49]. Future work could look into incorporating specific gene regulatory networks that mediate hypoxic cellular response in the context of cell migration and invasion [50]. Third, for sake of simplicity, we considered mutually exclusive phenotypes - ‘go- or-grow’, but recent preclinical and clinical work has emphasized the role of hybrid or intermediate phenotypes which can be the fittest for metastasis [51–53]. Finally, in numerical simulations described in this paper, we utilized parameter values from existing literature to demonstrate various qualitative features of the model variables. However, for a thorough parameter inference and estimation, further model simplification and enhanced spatially resolved experimental data will be beneficial [54, 55].

## 6 Appendices

### 6.1 Appendix A

Here, we drive the analytical solution of the proposed model in the spatial average sense for an oxygen-deprivation environment (*c < c*_*H*_). The model equations become

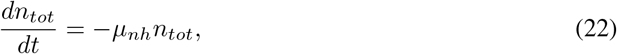

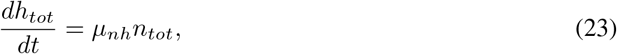

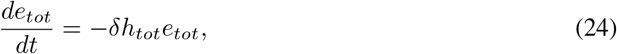

with the initial conditions *n*_*tot*_(*t* = 0) = *N*_0_, *h*_*tot*_(*t* = 0) = 0 and *e*_*tot*_ = *E*_0_. Here, *n*_*tot*_ = ∫_Ω_ *n*(*x, t*)*dx, h*_*tot*_ = ∫_Ω_ *H*(*x, t*)*dx, e*_*tot*_ = ∫_Ω_ *e*(*x, t*)*dx* and Ω = [0, *X*].

The solutions of Eqs. (22)-(24) are given as

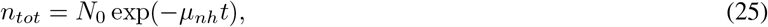

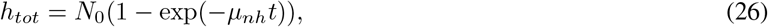

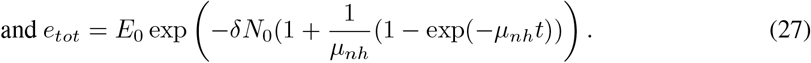

### 6.2 Appendix B

Here, we derive the solution of the spatially averaged model under oxygen-rich conditions, i.e., *c > c*_*H*_. In this environment, only normoxic cells proliferate. Since the tumor growth begins with a normoxic population, there is no hypoxic population (i.e., *h*_tot_ = 0), and the switching from hypoxic to normoxic is not activated. It is also noted that ECM is not degraded, as no hypoxic cells are present in the environment. Therefore, the total ECM density is a constant in this case i.e., *e*_*tot*_(*t*) = *e*_*tot*_. Hence, the spatially averaged model reduces to

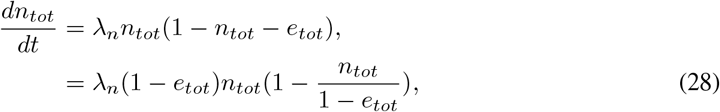

with the initial condition *n*_*tot*_ = *N*_0_. As *e*_*tot*_ is a constant, Eq. (28) becomes the standard logistic growth equation

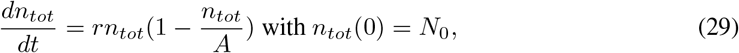

where *r* = *λ*_*n*_(1 −*e*_*tot*_) and *A* = (1 −*e*_*tot*_).

The solution of Eq. (29) is given as

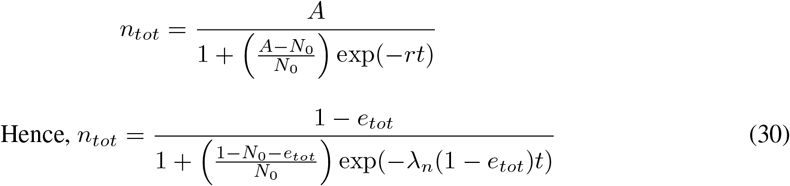

### 6.3 Appendix C

Here, we find out upper bound of timescale of memory induction and withdrawal. It is governed by

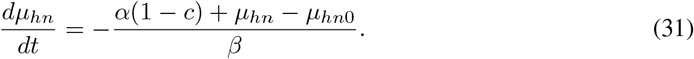

During normoxia i.e., *c* = 1, Eq. (31) becomes

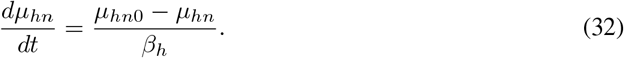

Its maximum value occurs when *μ*_*hn*_ = 0 and it is 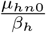. Therefore, the fasted timescale of memory erasing rate is 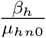. In our simulation, *μ*_*hn*0_ = 0.5 and 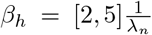. Hence, the timescales of memory dilution to range from four to ten cell divisions.

During hypoxia i.e., *c* = 0.4, Eq. (31) becomes

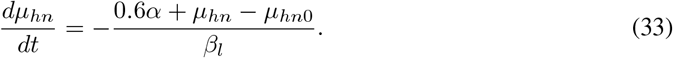

The maximum value of 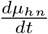 occurs when *μ*_*hn*_ = *μ*_*hn*0_ and the corresponding value is 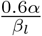. Hence, the fasted timescale of memory induction is 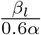. In our simulation, *α* = 0.8 and 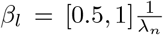. Hence, the timescale of memory induction is approximately half to two times of cell divisions.

## Data availability

Code used to produce the results are available on GitHub at https://github.com/gsadhuIITG/Hypoxic_memory

## Funding

JTG was supported by the Cancer Prevention and Research Institute of Texas (CPRIT RR210080) and the National Institute of General Medical Sciences of the National Institutes of Health (R35GM155458). JTG is a CPRIT Scholar in cancer research. MKJ was supported by Param Hansa Philanthropies.

## Ethical approval

The authors did not conduct any research with humans or animals.

## Conflict of interest

The authors declare no conflict of interest.

## Supplementary Information

**Fig. S1:**
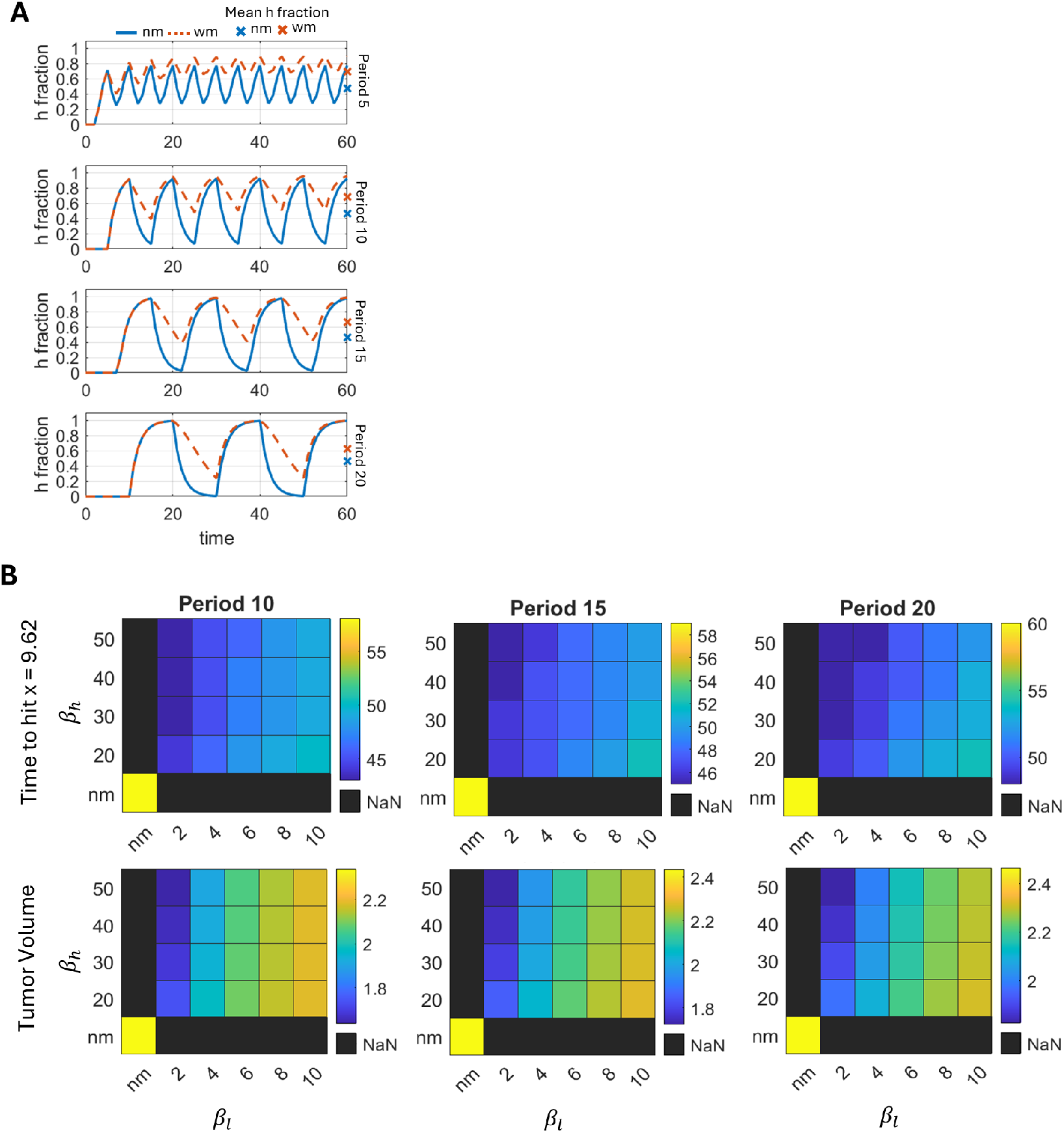
A) Temporal changes in the hypoxic cell fraction in the tumor bulk for increasing time period. ‘nm’ represents no hypoxic memory case and ‘wm’ denotes hypoxic memory with *β*_*h*_ = 20 and *β*_*h*_ = 2. Mean *h* fraction is calculated by averaging h fraction over the simulation period. B) Changes in the tumor volume and time to hit at a physical distance of x = 9.62 (minimum invasion length obtained for *β*_*h*_ = 20 and *β*_*l*_ = 10) for different combinations of memory induction, erasure timescales and environment period.

**Fig. S2:**
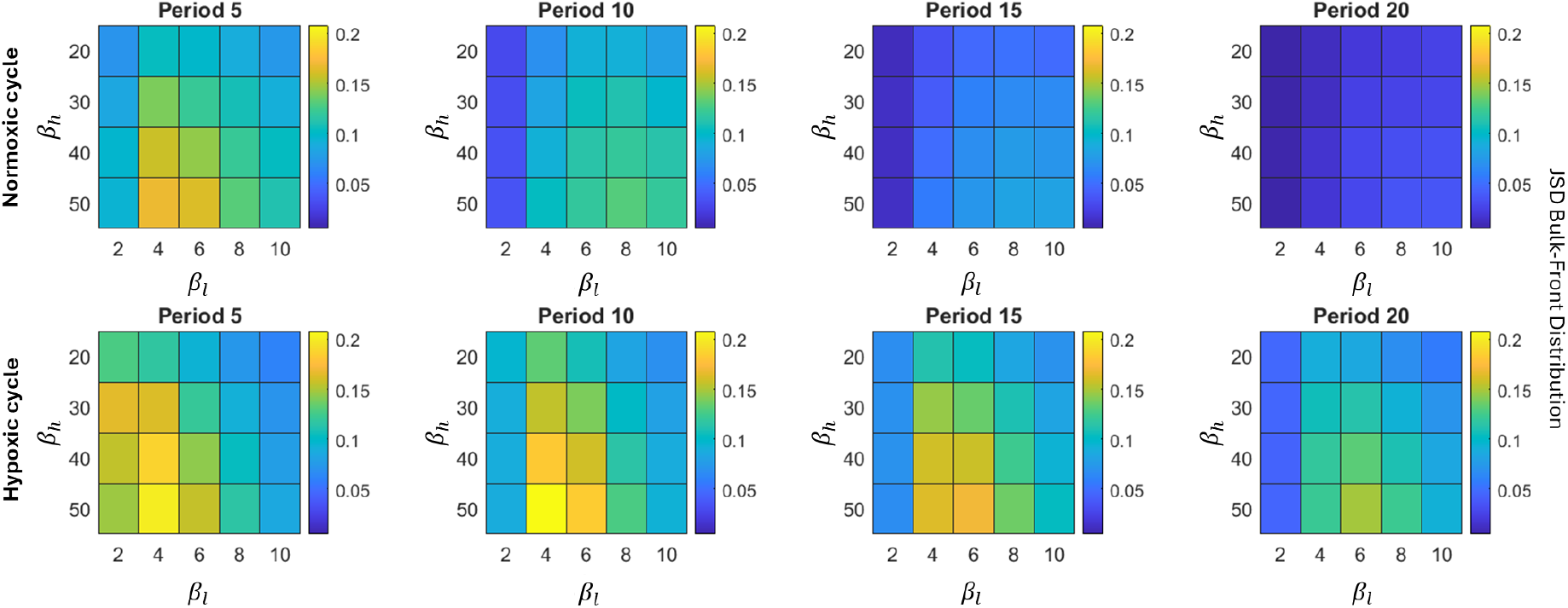
Using Jensen-Shannon Divergence to quantify the differences in phenotypic distributions of hypoxic cells between the tumor front and bulk cells at end of the normoxia and hypoxia environmental cycles. The hypoxia period bias is set to 0.5.

**Fig. S3:**
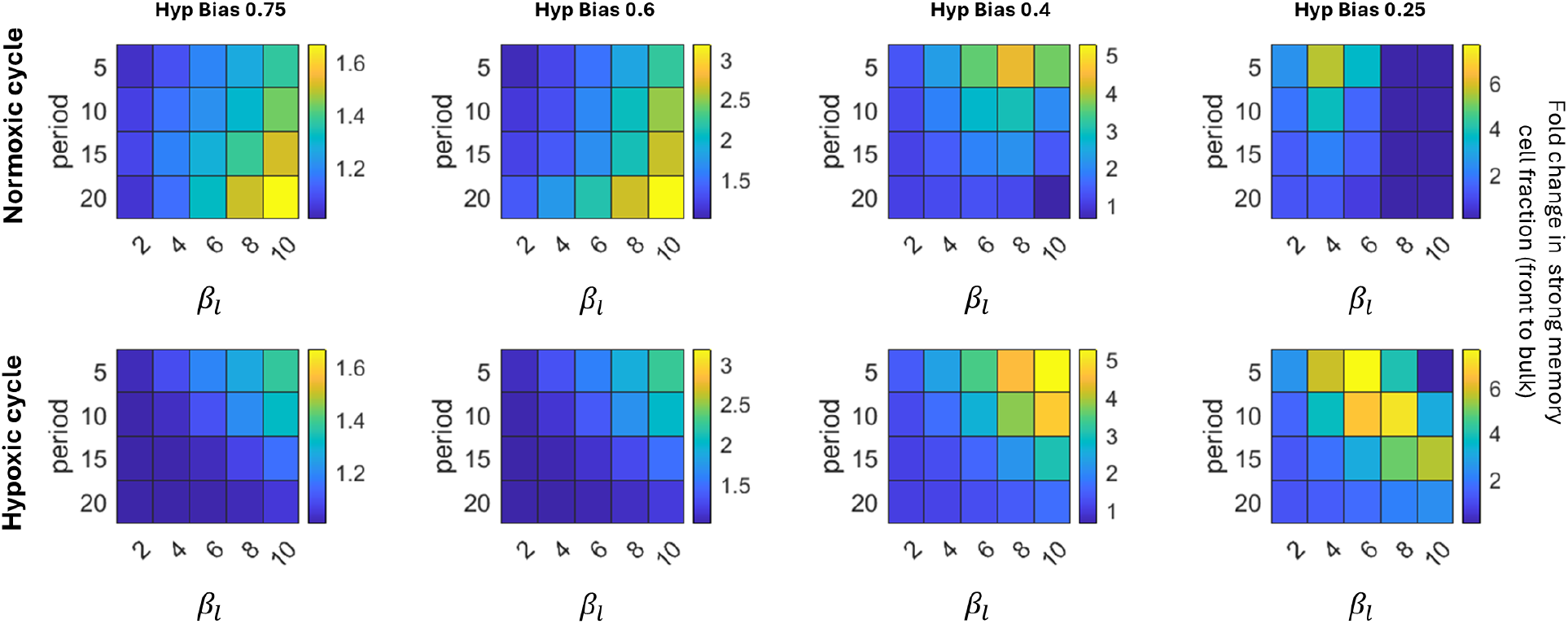
Quantification of ratio of strong memory cell fraction in the front to bulk regions at the end of normoxia and hypoxia cycles of different environmental bias. Here, the strong memory fraction is calculated by summing normalized cell density with *μ*_*hn*_ *<* 0.25. Results presented are with *β*_*h*_ = 20.

**Fig. S4:**
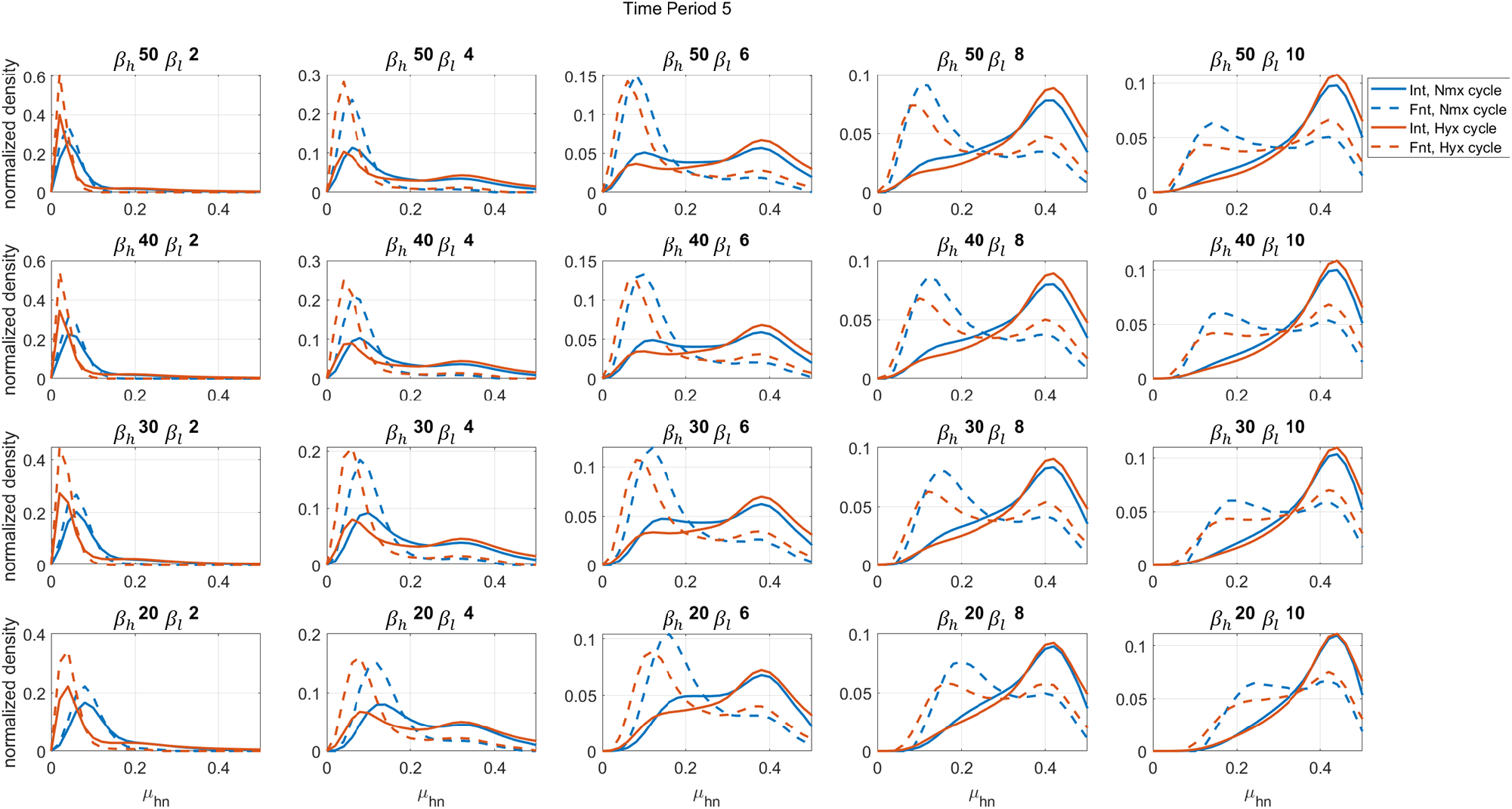
Normalised phenotypic (*μ*_*hn*_) distribution of cells in the bulk (total population) and front (defined by width of 0.5 unit distance from the right boundary) analysed at the end of the normoxic and hypoxic cycles of the last period of the simulation. Here the environment period is 5.

**Fig. S5:**
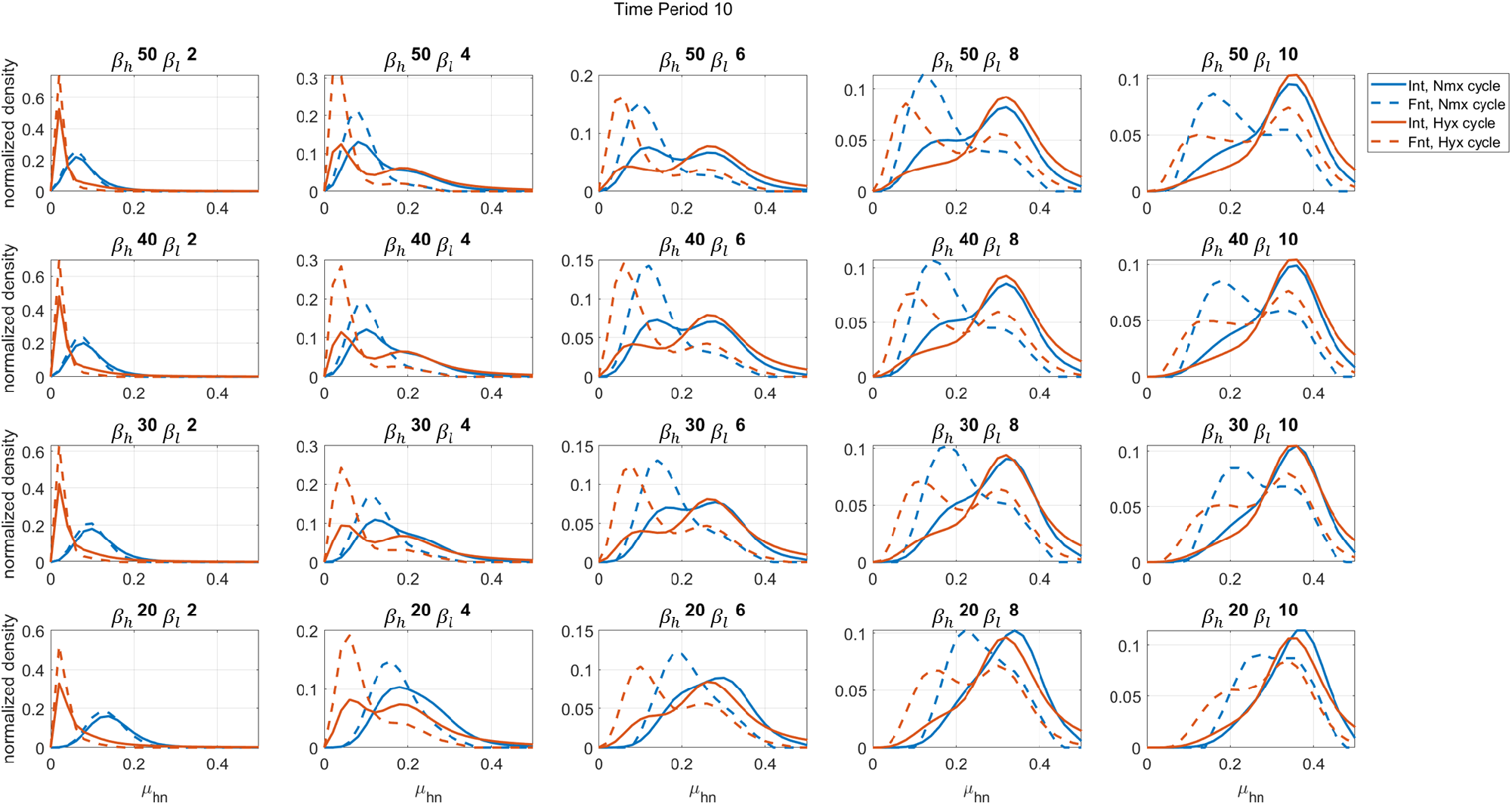
Normalised phenotypic (*μ*_*hn*_) distribution of cells in the bulk (total population) and front (defined by width of 0.5 unit distance from the right boundary) analysed at the end of the normoxic and hypoxic cycles of the last period of the simulation. Here the environment period is 10.

**Fig. S6:**
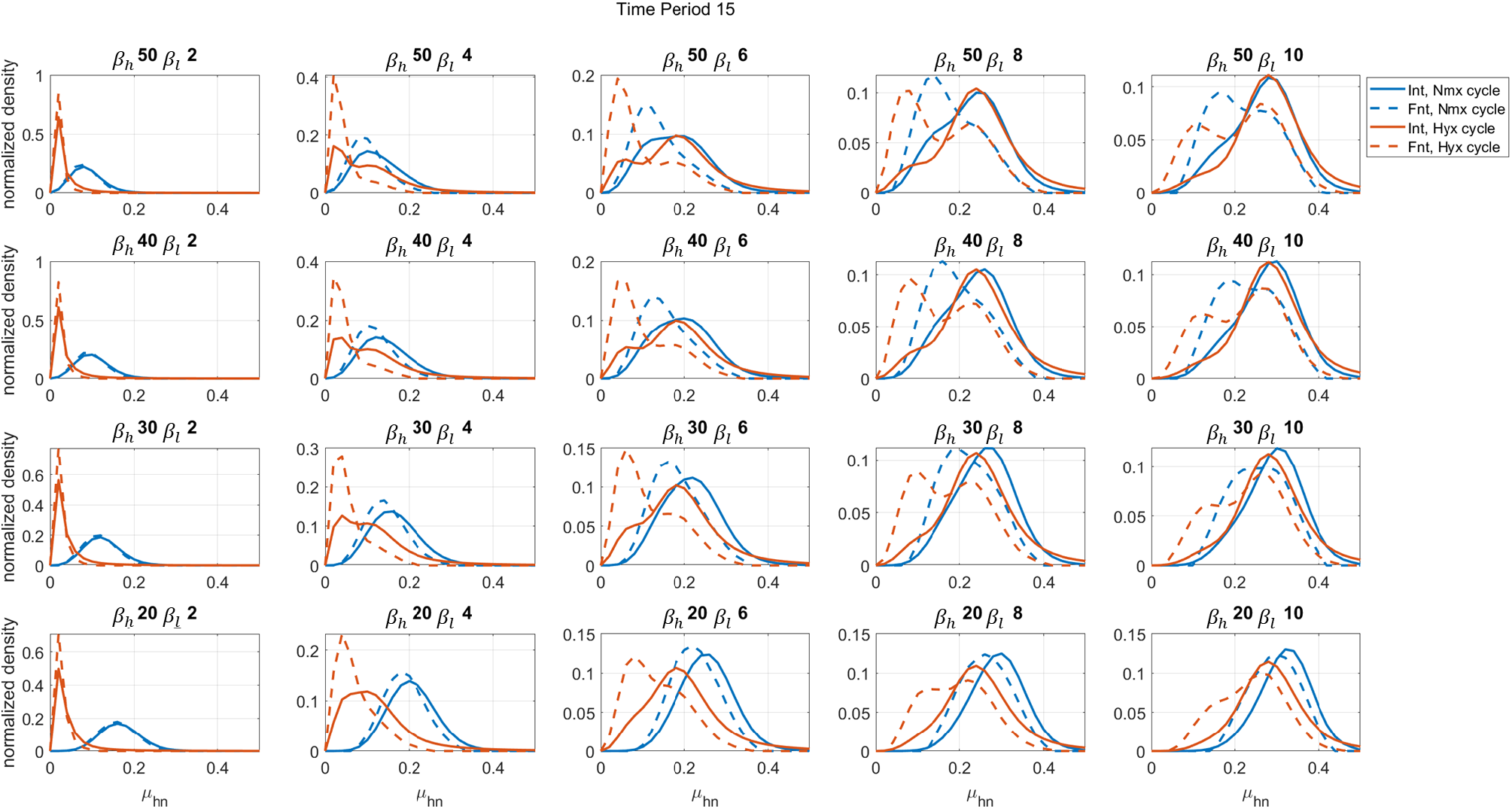
Normalised phenotypic (*μ*_*hn*_) distribution of cells in the bulk (total population) and front (defined by width of 0.5 unit distance from the right boundary) analysed at the end of the normoxic and hypoxic cycles of the last period of the simulation. Here the environment period is 15.

**Fig. S7:**
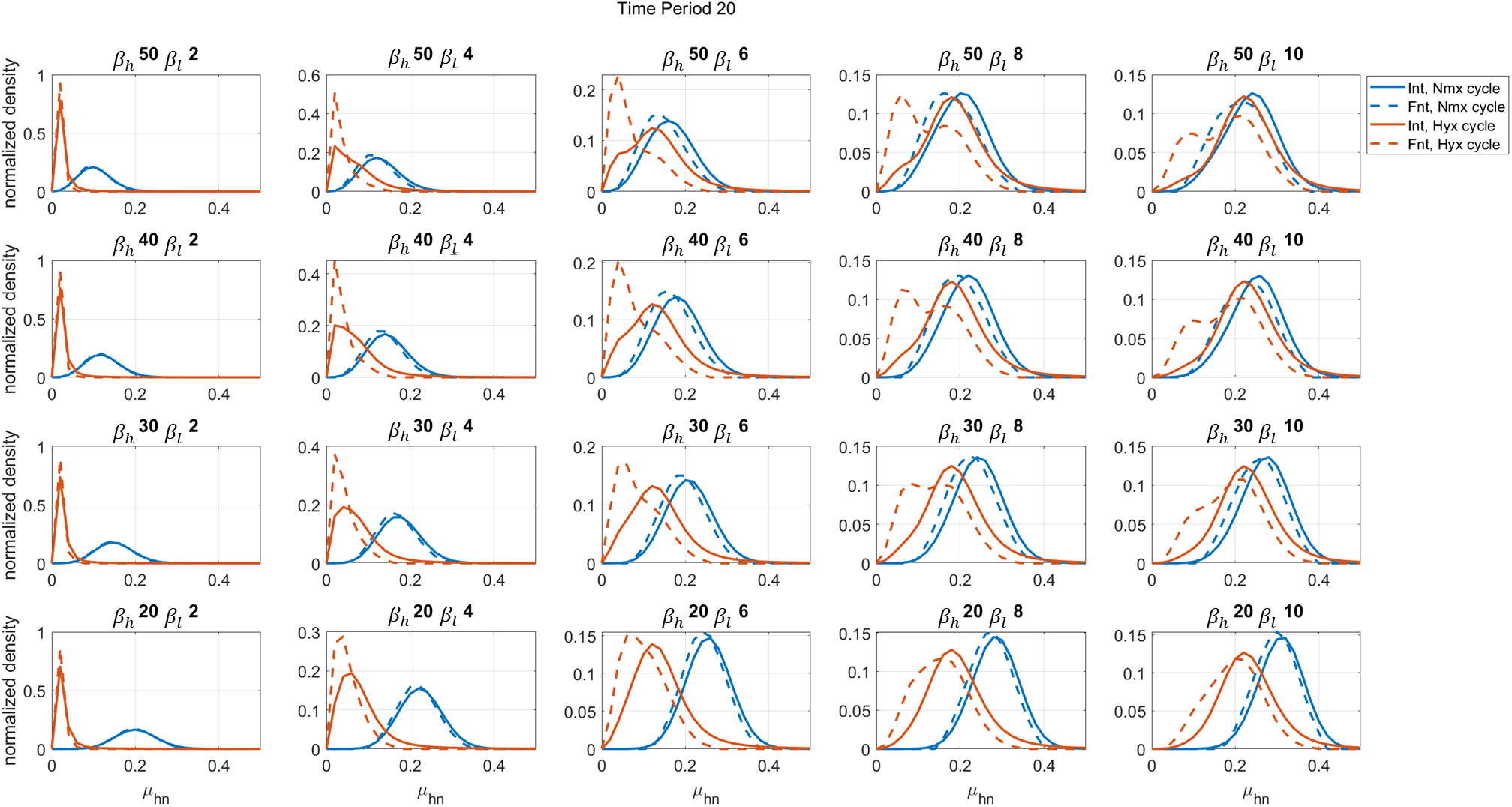
Normalised phenotypic (*μ*_*hn*_) distribution of cells in the bulk (total population) and front (defined by width of 0.5 unit distance from the right boundary) analysed at the end of the normoxic and hypoxic cycles of the last period of the simulation. Here the environment period is 20.

## Notes

### Competing Interest Statement

The authors have declared no competing interest.

